# A novel enhancer-AAV approach selectively targeting dentate granule cells

**DOI:** 10.1101/2023.02.03.527045

**Authors:** Emmie Banks, Claire-Anne Gutekunst, Geoffrey A. Vargish, Anna Eaton, Kenneth A. Pelkey, Chris J. McBain, James Q. Zheng, Viktor Janos Oláh, Matthew JM Rowan

## Abstract

The mammalian brain contains the most diverse array of cell types of any organ, including dozens of neuronal subtypes with distinct anatomical and functional characteristics. The brain leverages these neuron-type-specializations to perform diverse circuit operations and thus execute different behaviors properly. Through the use of Cre lines, access to specific neuron types has steadily improved over past decades. Despite their extraordinary utility, development and cross-breeding of Cre lines is time-consuming and expensive, presenting a significant barrier to entry for many investigators. Furthermore, cell-based therapeutics developed in Cre mice are not clinically translatable. Recently, several AAV vectors utilizing neuron-type-specific regulatory transcriptional sequences (enhancer-AAVs) were developed which overcome these limitations. Using a publicly available RNAseq dataset, we evaluated the potential of several candidate enhancers for neuron-type-specific targeting in the hippocampus. Here we identified a promising enhancer-AAV for targeting dentate granule cells and validated its selectivity in wild-type adult mice.

## Introduction

The hippocampus plays critical roles in pattern recognition, spatial navigation, and episodic memory. To accomplish these diverse tasks, a series of operations are executed in the ‘trisynaptic circuit’ (i.e., dentate gyrus [DG], CA3, and CA1), where information is functionally transformed by distinct operations at each location^1^. This is well illustrated by the highly divergent spatial coding schemes of distinct hippocampal subfields^2–4^ and their contributions to distinct aspects of mnemonic function^5^. However, due to the highly sequential manner in which information is processed through the trisynaptic circuit, the hippocampus is particularly susceptible to pathophysiology including epilepsy and various psychiatric disorders^6^. Therefore, subfield-specific cellular targeting methods are essential to understand the function of hippocampal areas in health and disease and to develop effective preventative and therapeutic interventions.

Our ability to investigate the function of hippocampal subfields in isolation has steadily improved in past decades. Researchers have long-exploited the anatomically well-separated nature of the subfields for lesion studies^7,8^. More recently, genetic targeting (i.e., Cre mice) approaches have become the standard for identification and discrimination between functionally heterogeneous cell populations. Cre lines with selectivity for specific excitatory cell types of each hippocampal subfield are available^9,10^. Despite their utility, Cre approaches for investigation of specific brain cell populations are nonetheless limited to studies in mice alone, are often expensive due to the need for extensive cross-breeding to other (floxed) mouse lines and cannot be directly translated to clinical therapies. Thus more versatile and translatable cell-specific vector targeting methods (enhancer-AAVs) are garnering increasing interest in neuroscience^11–13^. Enhancer-AAVs incorporate short transcriptional regulatory elements to target expression of transgenes selectively within distinct cell populations in wild-type animals *in vivo*, and often maintain cell-type-specificity across species including humans^12–14^. Despite a growing appreciation of their utility, enhancer-AAVs with selectivity for distinct hippocampal principal neurons are currently limited.

A recent study rigorously investigated differences in chromatin accessibility of single cell regulatory elements (scREs) across distinct neocortical pyramidal neurons, identifying several putative novel cell-type-specific enhancer elements. Based on these findings, enhancer-AAVs incorporating these scREs successfully targeted functionally distinct pyramidal cells residing in different cortical layers^14^. As an analogous approach in the hippocampus has not been established, we examined whether the scREs identified by Graybuck and colleagues discriminate between hippocampal excitatory neurons. By comparing the expression of these scRE-regulated genes with an established RNA-seq map of the hippocampus^10^, we identified a candidate enhancer exhibiting high selectivity for dorsal DG granule cells. When incorporated into an enhancer-AAV to drive YFP expression, we observed abundant and highly selective labeling of dentate gyrus granule cells in the dorsal hippocampus. Expression was present in all subcellular compartments of the granule cells, and importantly, no measurable expression was detected in other neighboring excitatory cell types in the CA3 or hilus. Together, we identified a novel viral approach to selectively label hippocampal granule cells, which should allow for specific presynaptic genetic modulation and targeted clinical interventions at the mossy fiber-CA3 synapse in the future.

## Results

### Identification of a DG granule cell-selective AAV targeting approach

Selective viral targeting of distinct excitatory cell classes in cortex and hippocampus is currently limited. The most widely used excitatory neuron promotor (i.e., CaMKII) induces transgene expression across most excitatory neuron classes in these regions. Furthermore, the CaMKII promoter also induces off-target transgene expression in inhibitory neurons^15^. Thus, identifying novel regulatory elements with specificity to functionally distinct excitatory neuron subclasses is essential. A recent study identified distinctions in chromatin accessibility among different cortical excitatory neurons within the same cortical region^14^, revealing several regulatory elements selective for distinct deep-layer pyramidal neurons. The hippocampus is also comprised of several neighboring excitatory neuron subtypes, which currently are not accessible using analogous single viral (i.e., enhancer-AAV) approaches.

Using a well-curated RNA-seq database from distinct hippocampal subregions^10^, we first examined whether four candidate genes from the Graybuck study (*Lratd2, Car3, Osr1, Hsd11b1*) displayed differential expression in hippocampal excitatory neuron subclasses in the mouse brain (**Figure 1A**). In cortical neurons, the *Lratd2, Car3, Osr1*, and *Hsd11b1* genes are associated with unique, mouse single-cell regulatory elements (mscREs) (mscRE4, mscRE10, mscRE13, and mscRE16, respectively)^14^. When cross-referenced with hippocampal neurons, we noted that three of these four genes were selectively expressed in distinct excitatory types (**Figure 1B**). The gene *Lratd2* (otherwise known as *Fam84b*), associated with the enhancer region mscRE4 (**Figure 1C**), was highly expressed in dorsal DG granule cells (**Figure 1B**, yellow bar), and to a lesser extent in ventral granule cells and dorsal CA1 pyramidal neurons, relative to other hippocampal neurons. The gene *Car3* displayed selective expression in ventral CA3 pyramidal neurons. The *Osr1* gene displayed relatively high expression in mossy cells and CA3 pyramids. Lastly, *Hsd11b1* showed relatively similar expression across all excitatory cells.

**Figure 1.**
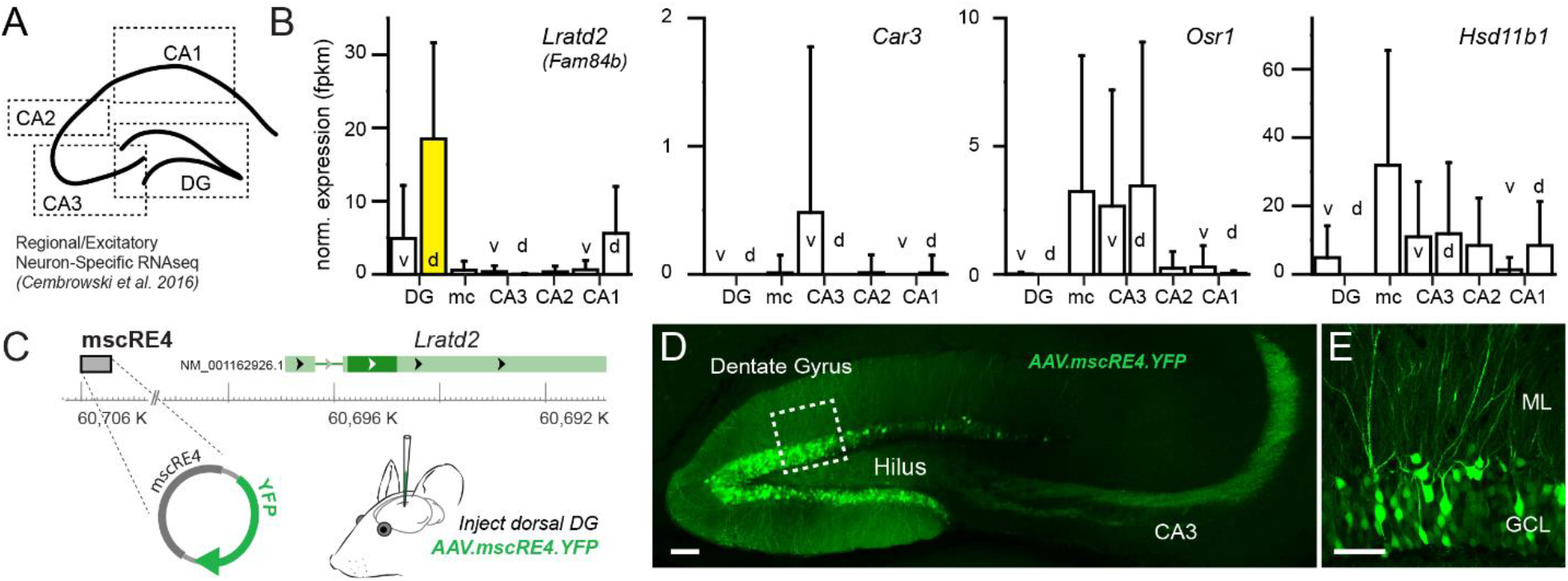
Identification of dentate granule cell-selective enhancer for AAV approach. (A) An RNA-seq database used to examine differential gene expression between all major excitatory neuron classes across the hippocampal subfields showed (B) *Lratd2* had high specificity to dentate gyrus (DG) granule cells in the dorsal DG and the highest overall normalized expression level compared to the other genes with associated enhancers previously shown to distinguish cortical excitatory neurons *in vivo*. (C) A plasmid containing three copies of a short core sequence of mscRE4 transcript were packaged into an AAV (PHP.eB) capsid for intracerebral injections, which resulted in (D, E) YFP expression in DG granule cells after 1-2 weeks expression. Scale bars: 100 µm (D) and 50 µm (E). *v = ventral, d = dorsal, DG = dentate gyrus, mc = mossy cells, CA3 = cornu Ammonis 3, CA2 = cornu Ammonis 2, CA1 = cornu Ammonis 1, ML = molecular layer, GCL = granule cell layer*.

If RNA expression levels (**Figure 1B**) correlate with chromatin availability of the associated mscREs in these hippocampal cell classes^16^, then AAVs that incorporate these enhancer sequences (enhancer-AAVs) might electively drive expression in distinct hippocampal cell classes. Because the mscRE4-associated *Lratd2* gene showed both the greatest overall expression and high specificity to a single excitatory cell class, we packaged a plasmid (Addgene# 164458) containing three copies of a short “core” mscRE4 enhancer sequence to express a YFP reporter into the PHP.eB capsid^17^ (referred to as AAV.mscRE4.YFP hereafter) (**Figure 1C**). ∼150 nl of the AAV.mscRE4.YFP virus was stereotactically injected into the dorsal DG of mature C57Bl/6J mice. In accordance with RNA expression of the mscRE4-associated *Lratd2* gene (**Figure 1B**), DG granule cells and their mossy fiber axons displayed abundant YFP labeling (**Figure 1D, E**) 1-2 weeks post-injection. Remarkably, somatic YFP expression in regions near the granule cell layer (i.e., the hilus and CA3 regions) was absent (**Figure 1D**). We also evaluated whether AAV.mscRE4.YFP selectively targets pyramidal tract (i.e., layer 5B) neurons in cortex, as previously reported in the primary visual area^14^, using stereotactic injections in the primary motor cortex. As anticipated, we observed highly selective expression in M1 layer 5B pyramidal neurons (**Supplemental Figure 1**). Together, our results indicate that mscRE4-based enhancer-AAVs are well suited for granule cell targeting with local stereotactic injections.

### AAV.mscRE4.YFP labels DG granule cells but not neighboring hilar mossy cells

Our transcriptomic and histological surveys suggest that the mscRE4 enhancer preferentially drives expression in granule cells with respect to neighboring excitatory neurons in the dorsal hippocampus. The CaMKII promoter has traditionally been incorporated in AAVs to target excitatory neuron types. However, CaMKII promotes reporter expression indiscriminately across all major excitatory cell classes in close proximity to the DG, including granule cells, hilar mossy cells, and pyramidal neurons^15,18,19^. Thus, we examined the co-expression patterns of AAV.mscRE4.YFP and AAV.CaMKII.mCherry (Addgene# 114469) from mice unilaterally co-injected with both AAVs (> 1^13^ vg/ml each) in the dorsal hippocampus. To control for potential differences in serotype tropism^20^, the AAV.CaMKII.mCherry construct was also packaged into the PHP.eB capsid. 1-2 weeks post-injection in dorsal DG, slices were prepared using a derivative of the ‘magic cut’^21^ to preserve granule cell mossy fiber axons and mounted for confocal imaging (**Figure 2A**). mCherry labeling was consistently observed in granule cells somas and their dendrites in the DG, pyramidal cells in the CA3 stratum pyramidale, and in putative hilar mossy cells. However, co-labeling of YFP and mCherry was highly selective for granule cells (**Figure 2B**) with no apparent co-labeling of somas in the hilus. Specifically, hilar cells positive for mCherry were consistently negative for YFP despite their proximity to the injection site (**Figure 2C**). In contrast, 77.2% of mCherry-positive cells in the DG granule cell layer were also YFP-positive, indicating relatively strong transgene expression directed by mscRE4 in these cells (**Figure 2C**). Importantly, AAV.CaMKII.mCherry labeling extended to neurons well outside of the DG, indicating that spatial restriction of AAV diffusion from our injection site was unlikely to account for the granule cell selectivity of mscRE4.

**Figure 2.**
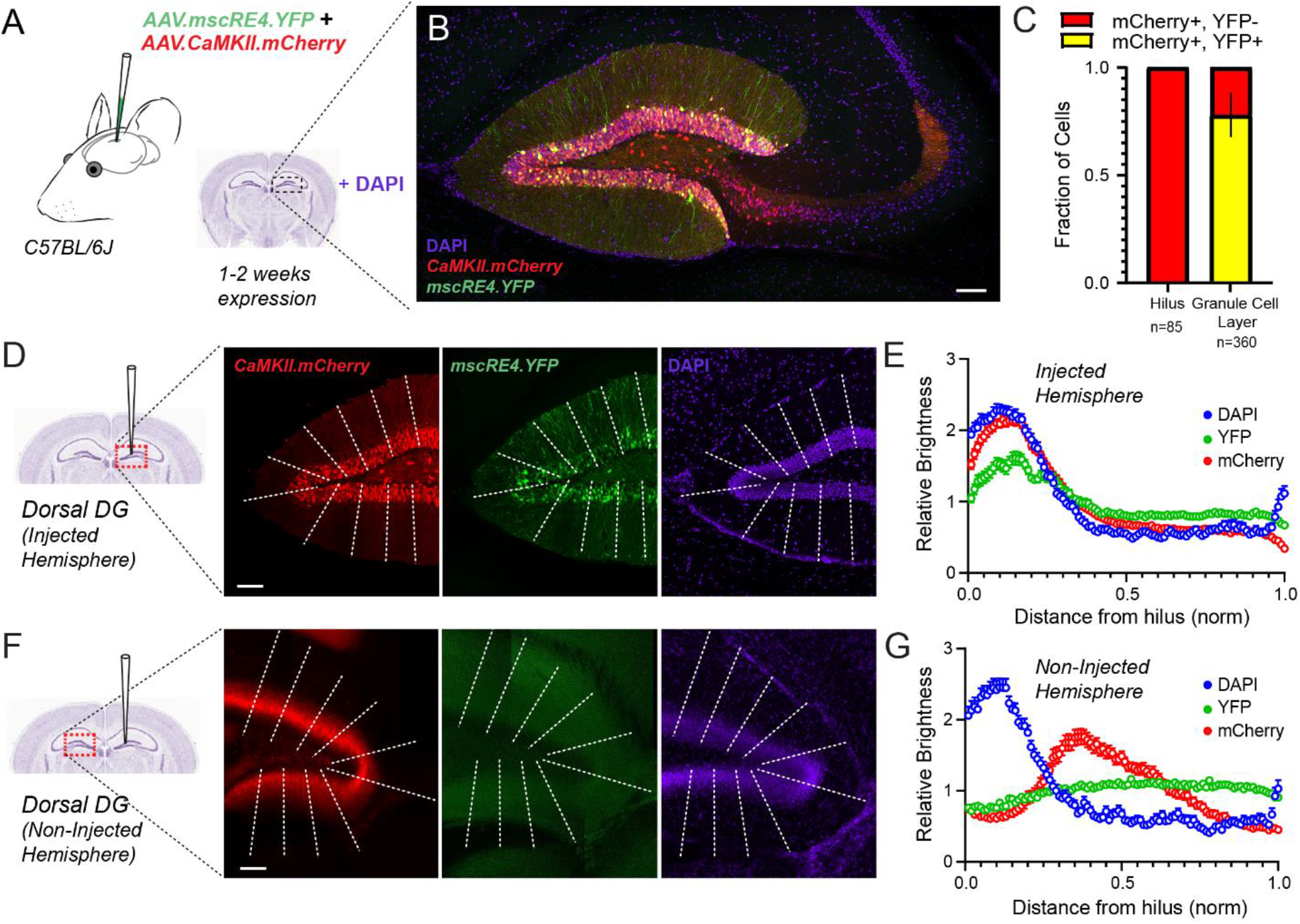
AAV.mscRE4.YFP labels dentate granule cells in dorsal dentate gyrus. (A) AAV.mscRE4.YFP and AAV.CaMKII.mCherry were unilaterally injected together in the dorsal dentate gyrus (DG) in one hemisphere of young adult (7-11 week old) wild-type mice (n = 5; 3 females and 2 males), and expressed for 1-2 weeks prior to acute slice preparation. (B) YFP/mCherry co-expression appeared selective for granule cells in confocal images (40x objective; oil) of ‘magic cut’ slices, which was further assessed by (C) cell counting in the DG-hilus region, where mCherry+ hilar cells were consistently YFP- and 77.2% of mCherry+ cells in granule cell layer were YFP+. (D, E) Line scan profiles in the DG in the injected hemisphere and (F, G) in the non-injected hemisphere depicting the relative brightness of DAPI, YFP, and mCherry signals across a normalized distance from the hilus. Scale bars: 100 µm throughout

To further examine the neuron-type specificity of AAV.mscRE4.YFP within DG-hilus region, we next quantified the relative brightness of YFP, mCherry, and DAPI signals in the DG of the injected hemisphere as well as the non-injected hemispheres (**Figure 2D, 2F**) of the same mice. In both injected and non-injected hemispheres, the peak DAPI signal delineated the granule cell layer in the DG (**Figure 2E, 2G**). In injected hemispheres, mCherry and YFP signals followed the DAPI signal, rapidly tapering by around 50% (**Figure 2E**), supporting our claim that AAV.mscRE4.YFP selectively labels granule cell bodies and their neurites.

Hilar mossy cell axons make strong contralateral projections, terminating in the inner molecular layer of the contralateral dentate gyrus^22^. Thus we also examined the labeling via mCherry and YFP in the non-injected DG (**Figure 2F; Supplemental Figure 2**). In non-injected hemispheres, the mCherry signal was well structured in the DG but highly disjointed from DAPI (**Figure 2G**), signifying AAV.CaMKII.mCherry labeling of commissural hilar mossy cell axons. The mscRE4-driven YFP signal was notably flat in non-injected hemispheres (**Figure 2F, G**), providing further evidence for mscRE4 as a granule cell-selective enhancer that spares mossy cells.

### AAV.mscRE4.YFP labels mossy fiber axons but not neighboring CA3 pyramidal neurons

We next evaluated expression in the CA3 region following co-injection of AAV.CaMKII.mCherry and AAV.mscRE4.YFP in the dorsal DG. Granule cells make direct projections to CA3 via mossy fibers (**Figure 3A**), which were preserved in our slices using the ‘magic cut’ (see Methods). We observed strong somatic expression of mCherry in both cell somas and neuritic structures of CA3 (**Figure 3B**), likely due to mCherry expression in both mossy fiber axons and CA3 pyramidal cells. As described above, strong YFP expression was present in the mossy fibers extending into CA3. However, in contrast to the mCherry signal, expression of YFP was absent from cell somas in CA3 (**Figure 3B**). Furthermore, YFP expression was absent from mCherry-positive CA3 pyramidal neurons surrounded by co-labeled mossy fiber axons projecting from the DG (**Figure 3C1-3C3 & Supplemental Figure 3;** examples from 3 different mice). Combined, our findings indicate high selectivity of the mscRE4 enhancer-AAV for granule cells over neighboring excitatory cell types.

**Figure 3.**
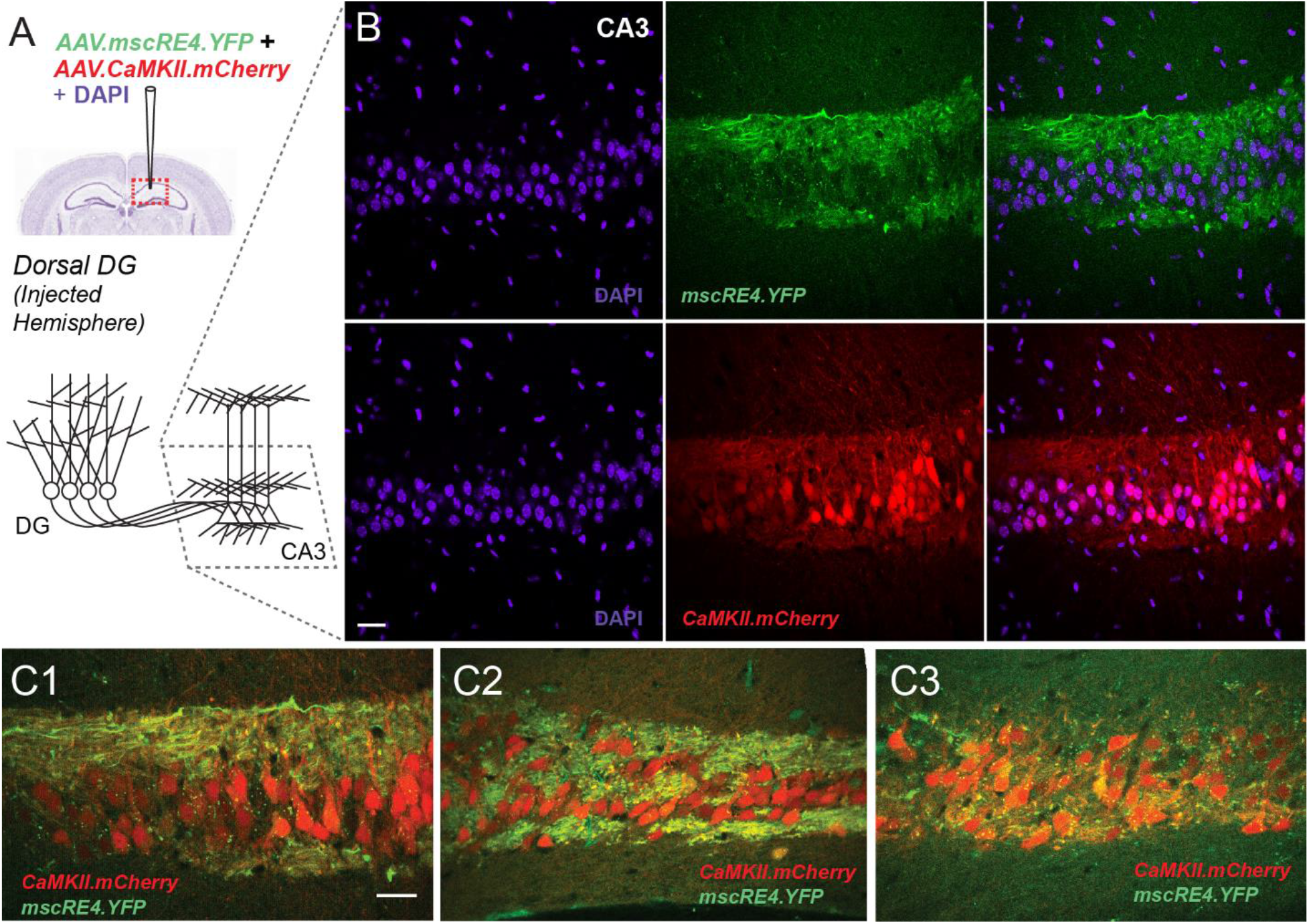
AAV.mscRE4.YFP expression in mossy fibers but not CA3 pyramidal neurons. (A) Co-injection of AAV.mscRE4.YFP and AAV.CaMKII.mCherry into the dorsal dentate gyrus (DG) in one hemisphere of adult (7-11 weeks old) wild-type mice produced (B) mCherry expression in CA3 somas along with a lack of YFP expression in CA3 somas, as well as (C1-C3) co-expression of mCherry and YFP in mossy fiber axons projecting from the DG (as seen in confocal images; 40x objective, oil). Scale bars: 50 µm

A recent study found that anterograde AAV serotypes (i.e., PHP.eB) may induce dose-dependent toxicity specifically in adult-born granule cells.^34^ This problem could be circumvented through use of AAV-retro capsids to direct expression in granule cells. As DG granule cells make direct projections to CA3, we sought to determine whether a retrograde form of AAV.mscRE4.YFP would show a similar targeting profile when injected in the CA3 region. Thus, we packaged the same SYFP-expressing mscRE4 construct into a retrograde capsid (referred to as AAV.retro.mscRE4.YFP hereafter) and injected it at the CA3b/CA3a border (**Supplemental Figure 4A**). After 1-2 weeks, we observed a YFP expression pattern comparable to that produced by the anterograde (PHP.eB) AAV.mscRE4.YFP capsid. Specifically, in the DG we observed YFP expression in granular layer cell somas and dendrites in the molecular layer (**Supplemental Figure 4B**), while in CA3, punctate YFP expression surrounding but not overlapping with pyramidal cell somas was again apparent (**Supplemental Figure 4C**). Together this indicates that AAV.retro.mscRE4.YFP injections in the CA3 region target DG granule cells via their mossy fiber axons. The density of YFP expression in the DG granular layer appeared reduced compared to that produced by AAV.mscRE4.YFP, likely due to the different serotype and injection site. To confirm the extent and cell-type selectivity of AAV.retro.mscRE4.YFP expression using another method, we also performed immunohistochemistry to visualize expression with diaminobenzidine (DAB). We observed abundant dark brown reaction products in DG granular layer cell somas and molecular layer dendrites (**Supplemental Figure 4D**), as well as punctate expression in the CA3 injection region (**Supplemental Figure 4E**), thus complementing our AAV.retro.mscRE4.YFP expression seen with fluorescent confocal imaging.

**Figure 4.**
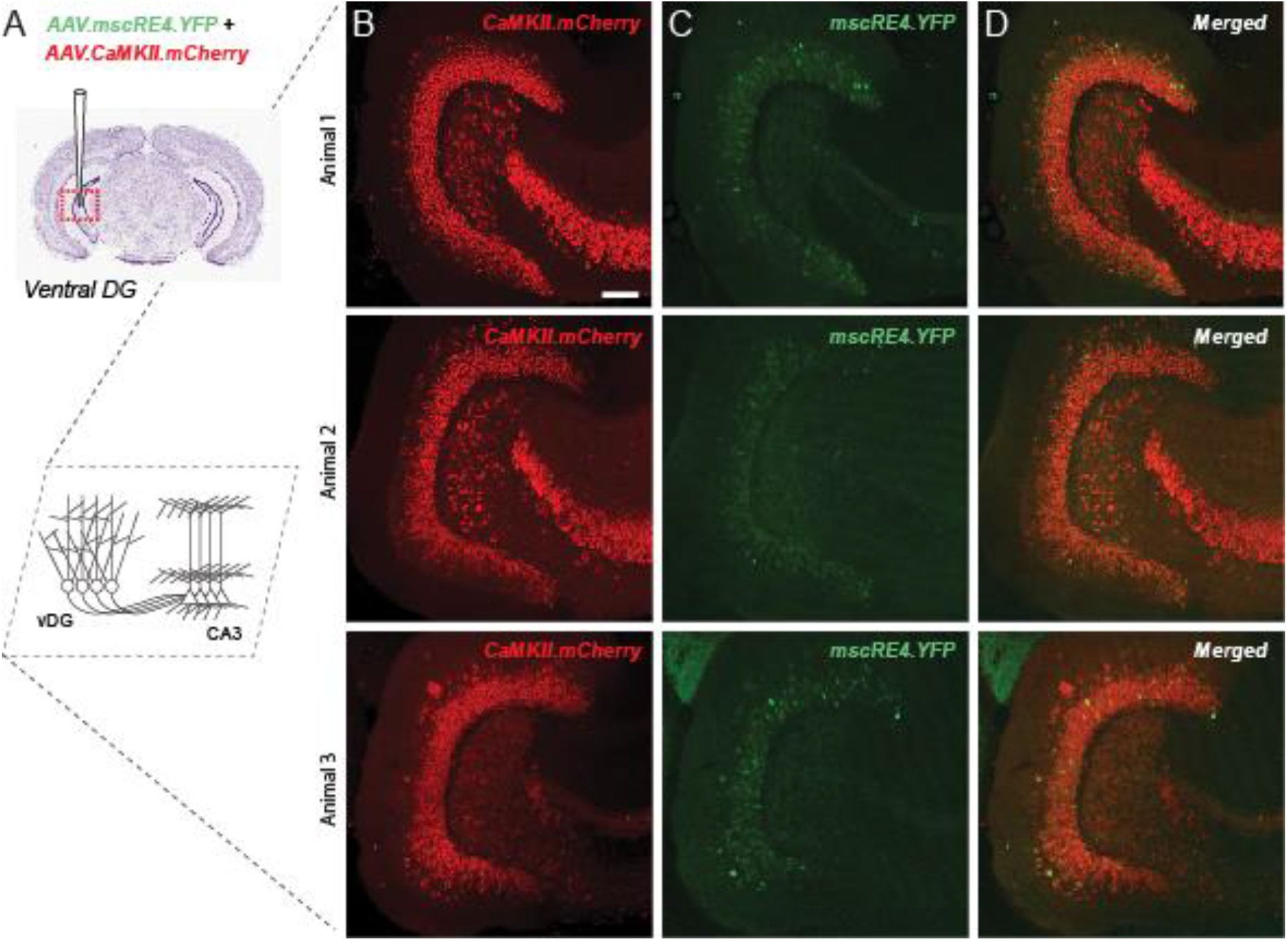
AAV.mscRE4.YFP and AAV.CaMKII.mCherry labeling in ventral hippocampus. (A) Adult wild-type mice were co-injected with AAV.mscRE4.YFP and AAV.CaMKII.mCherry in the ventral dentate gyrus (DG). After 1-2 weeks, in confocal images (20x objective; horizontal sections) we observed (B) mCherry labeling in DG granule cell somas, as well as putative hilar mossy cells and CA3 pyramidal cells. We also observed (C) YFP labeling in a subset of DG granule cells somas, with a noticeable absence of somatic labeling in the hilus or CA3, and thus (D) co-expression of mCherry and YFP labeling was restricted to the DG granular layer. Scale bars: 100 µm. Supplemental Figure 1.

### AAV.mscRE4.YFP labeling in ventral hippocampus

The mscRE4-associated *Lratd2* gene also showed expression in ventral DG granule cells (**Figure 1B**), albeit far less than that observed in the dorsal DG. Thus we also examined the expression patterns of AAV.mscRE4.YFP and AAV.CaMKII.mCherry from mice co-injected with both AAVs in the ventral DG (**Figure 4A**). As seen in the dorsal hippocampus, we observed mCherry labeling in DG granule cell somas, as well as putative hilar mossy cells and CA3 pyramidal cells (**Figure 4B**). We also observed YFP labeling in a subset of DG granule cells somas, with a noticeable absence of somatic labeling in the hilus or CA3 (**Figure 4C**). The overlap in mCherry and YFP labeling was again restricted to the DG granular layer in ventral hippocampus (**Figure 4D**). These findings were again in concordance with predictions from the RNAseq dataset (**Figure 1B**), as the density of YFP expression in the ventral DG granular layer appeared reduced compared to that produced in the dorsal hippocampus.

### Non-granule cell ‘off-target’ mscRE4 labeling caveats in dorsal hippocampus

While mscRE4-directed expression was primarily restricted to granule cells following local injection in dorsal DG, when injections were improperly targeted, we also noted dense YFP expression in the dorsal subiculum (**Supplemental Figure 5**). Furthermore, as anticipated by our RNAseq analysis earlier, we also saw sparse YFP expression in the putative CA1 pyramidal neurons when injections spread into that region (**Supplemental Figure 5**). In sum, these data indicate that mscRE4 is selective but not completely specific for granule cells in the dorsal hippocampal region.

## Discussion

Here we first examined the likelihood that regulatory elements with specificity for neocortical excitatory neuron subtypes would also map onto subsets of excitatory neurons in the hippocampus in wild type mice. Predictive analysis was performed by comparing hippocampal excitatory cell-specific mRNA expression for genes^10^ associated with newly identified regulatory regions of interest^14^. Several candidates with potential selectivity for different excitatory neurons were identified. A single promising candidate (mscRE4), which displayed high selectivity for dentate granule cells in dorsal hippocampus, was validated via local injection. To our knowledge, this is the first vector targeting method suitable for a single excitatory cell type within the hippocampal trisynaptic circuit. Our results suggest that highly distinct neuron types situated in different brain regions may share a similar chromatin accessibility signature and thus, effective enhancer elements. Thus, high-resolution comparisons of enhancer-viruses between these and other brain regions may advance cell-specific targeting technologies more rapidly.

### Cell-type-specific targeting with AAVs in the brain

Cre mouse lines are beneficial for cell-specific studies throughout the brain^10,23–25^; however, they are often limited by off-target^26–28^ or incomplete^29^ expression. The use of AAV vectors with cell-type-selective *cis*-active DNA control elements has opened an alternative strategy avenue to advance circuit, cellular, and synapse-specific questions in neuroscience. Inclusion of full-size or truncated promotor sequences can direct AAV expression within particular neuron classes^30–32^. However, the large size of most promotor sequences represents a significant issue due to the relatively small AAV packaging size. Shorter regulatory (i.e., enhancers) elements present an ideal opportunity for inclusion in AAV-targeting experiments, with several recent studies demonstrating exquisite neuron-subclass specificity^12,33^. An additional benefit of enhancer-AAVs is their cross-species adaptability, with specific enhancer elements neuron-type-specificity across rodents, primates, and mature human neurons^12,13^.

### mscRE4-driven expression in dorsal hippocampus

Recent work from the Allen Institute identified several short regulatory elements with specificity for distinct neocortical pyramidal cells, which were then validated in the V1 region after incorporation into enhancer-AAVs^14^. While many of the genes regulated by these newly identified enhancers showed interesting expression patterns in hippocampus^10^, here we focused on one candidate (mscRE4) showing selective and relatively high expression in excitatory granule cells in the dentate gyrus of the dorsal hippocampus. When injected locally, we found that mscRE4-directed YFP expression in granule cells, their dendritic in the dentate, and broadly throughout their mossy fiber axons. Intriguingly, there was an evident lack of expression in the nearby excitatory mossy cells and CA3 pyramidal neurons, matching our expectations based on the expression pattern of the *Lratd2* mRNA expression. Further, as our RNAseq results suggested, mscRE4-directed YFP expression was present, albeit to a much lower extent, in ventral hippocampal DG granule cells. Importantly, recent work suggests that adult-born granule cells may be susceptible to AAV-induced toxicity in a dose-dependent manner following direct injections into the dentate^34^. Thus viral titer may be a critical variable when studying granule cells with enhancer-AAVs. Alternatively, mscRE4 constructs may be packaged into the AAV-retro capsid and injected into CA3 to avoid potential toxicity in adult-born cells^34^.

It is unclear if a similar expression pattern will emerge in the dorsal hippocampus with systemic (i.e., via retro-orbital injection) introduction of this PHP.eB serotype^17^. However systemic injections with AAV.mscRE4.YFP will not be isolated to hippocampus, as highly specific expression in L5 cortical pyramidal cells has already been well demonstrated by the Allen Institute studies. Indeed, we also observed specific expression in L5B neurons when AAV.mscRE4.YFP was injected locally in M1 cortex, extending the use of mscRE4 in cortical L5 neurons to this cortical region as well. AAV.mscRE4 also appears to provide excellent utility for directed studies of the subiculum, where it strongly labeled putative excitatory neurons in this region. As with studies in dentate described here, accurately positioned, low-volume viral injections will undoubtedly be necessary for subiculum-specific targeting. Finally, YFP expression was occasionally seen in putative CA1 pyramidal neurons. While this ‘off-target’ effect does lessen the specificity of mscRE4 in granule cell studies, it could provide an avenue for experiments requiring highly sparse expression in CA1 in mice and potentially across other species.

### Basic science and clinical applications for granule cell targeting in dorsal hippocampus

While our findings indicate mscRE4-directed expression selectively in granule cells, additional studies employing mscRE4-driven expression of neuronal effectors (i.e., optogenetics) need to be performed in the same context to validate its specificity more thoroughly. Control of mossy fiber axon firing using ChR2 has been shown using a granule cell-targeting Cre line^35^. mscRE4 should allow for viral deployment of ChR2, genetically-encoded Ca2+ indicators, and any of their derivatives *in vivo* across any mouse line. Thus this enhancer virus has the potential to rapidly accelerate research into dentate-CA3 related questions, such as mechanisms of memory formation and storage, pattern separation, exploratory behaviors^5,36–38^, and similar questions in primates^39^ if cross-species expression is maintained.

Beyond basic science applications, mscRE4-granule cell targeting may have important preclinical applications. One of the prime advantages of cell-specific targeting with enhancer-AAVs is their rapid deployment in mouse models of different brain disorders^40^. Relevant to our findings, recent work using a multiplexed Cre approach demonstrated online control of epileptic seizures through optogenetic modulation of dentate gyrus excitability *in vivo*^41^. Thus, temporally well-controlled opto- or chemogenetic modulation of granule cell excitability with mscRE4 targeting represents a promising future direction for translational research in epilepsy and other disorders.

## Acknowledgments

We thank Ken Pelkey (NIH-NICHD) for his critical reading and helpful comments on the original manuscript draft and Annie Goettemoeller (Emory GDDBS) for help in surgical training supporting these studies.

## Funding

The current work was supported by these grants:

1R56AG072473 (MJMR)

1RF1AG079269 (MJMR)

## Methods

### Intracranial viral injections

All animal procedures were approved by the Emory University IACUC. Young adult (7-11 week old) C57Bl/6J mice of both sexes were used for all experiments, with data collected from ≥ 3 mice per experimental condition. One group of mice was initially injected with AAV(PHP.eB).mscRE4.minCMV.SYFP2 (AAV.mscRE4.YFP; titer 1.90E+13 vg/mL) alone, while another group was injected simultaneously with both AAV.mscRE4.YFP and AAV(PHP.eB).CaMKIIα.mCherry (AAV.CaMKII.mCherry; titer 6.00E+14 vg/mL) at a 1:1 ratio in the dentate gyrus region of the dorsal hippocampus. A final group of mice were injected with AAV(retro).mscRE4.minCMV.SYFP2 (AAV.retro.mscRE4.YFP; titer 7.36E+11 vg/mL) in the CA3b/CA3a border region of the dorsal hippocampus. When performing viral microinjections, mice were head-fixed in a stereotactic platform (David Kopf Instruments) using ear bars while under isoflurane anesthesia (1.8 - 2.2%). Thermoregulation was provided by a heating plate using a rectal thermocouple for biofeedback, thus maintaining core body temperature near 37°C. A craniotomy was gently made in the skull (< 0.5 μm in diameter) to allow access for a glass microinjection pipette. For dorsal dentate gyrus targeting, coordinates (in mm from Bregma) for AAV.mscRE4.YFP microinjection were X = +/−1.50; Y = −1.91; α = 0°; Z = −2.20. For ventral dentate gyrus targeting, coordinates were X = +/−2.61; Y = −3.52; α = 0°; Z = −3.65, −3.35 (virus delivered at two Z depths). For dorsal CA3b/CA3a border targeting, coordinates for AAV.retro.mscRE4.YFP microinjection were X = +/−2.50; Y = −2.00, α = 0°, Z = −2.25. For cortical M1 injection, coordinates were X = -1.75; Y = 1.41; α = 0°; Z = -0.850 and -0.55. Virus was injected slowly (∼0.02 μL min^-1^) using a Picospritzer (for dentate, 0.15-0.2 μl; for M1, ∼0.25 μl for each Z injection depth). After ejection of virus, micropipettes were held in place (≥ 5 min) before withdrawal. The scalp was closed with surgical sutures and secured with Vetbond (3M) tissue adhesive. After allowing for onset of expression (hippocampus, 1-2 weeks; cortical, 3 weeks), animals were sacrificed and acute slices harvested.

### ‘Magic cut’ slice preparation and tissue fixation

Mice were anesthetized with isoflurane and sacrificed by decapitation. The brain was immediately removed by dissection and submerged in ice-cold cutting solution (in mM) 87 NaCl, 25 NaHO3, 2.5 KCl, 1.25 NaH2PO4, 7 MgCl2, 0.5 CaCl2, 10 glucose, and 7 sucrose. Brain slices (100-200 μm) were sectioned using a vibrating blade microtome (VT1200S, Leica Biosystems) in the same solution. The ‘magic cut’^21^ was employed to maintain the GC-CA3 mossy fibers within the slice. To obtain the magic cut, following dissection from the cranium brains were cut with a scalpel to remove the cerebellum (*α = 90°)* and the prefrontal cortex (*α =90°)*. A medial cut was then made straight through the midline (*α = 90°)*. Each hemisphere then received a cut angled ∼22*°* inward to the midline at the anterior side of the brain. Acute slices were then transferred to a well-plate to fix in 4% paraformaldehyde (in PB) for 1-1.5 hours at room temperature. After fixation, slices were washed in 1xPBS for 5 minutes. A subset of slices were then incubated in DAPI in 1xPBS (1:1000) for 5 minutes. Slices were then mounted onto glass slides with Slowfade Diamond Antifade (Invitrogen), cover-slipped, and stored at 4°C until imaging.

### Immunohistochemistry

For a subset of slices obtained from mice injected with AAV.retro.mscRE4.YFP, immunohistochemistry was performed to further assess viral expression. Free-floating sections from one series were incubated in 0.1% (v/v) TritonX-100 and 3% (w/v) hydrogen peroxide to eliminate endogenous peroxidase, rinsed in PBS, and pre-blocked in 5% (v/v) normal donkey serum for 30 minutes at room temperature (RT). Sections were incubated in rabbit anti-GFP antibody (1:1000, A111222, Life Technologies, Carlsbad, CA) in PBS at RT for 24 hours, then rinsed and incubated for 1 hour at RT in biotinylated donkey anti-rabbit antibody (1:250, 711-065-152, Jackson Immunoresearch Laboratories, West Grove, PA) in PBS. After several rinses in PBS, the sections were incubated in avidin–biotin complex (ABC Elite; Vector Laboratories, Burlingame, CA, USA) for 90 minutes at 4 ºC. Immunoreactivity was visualized by incubation in 0.05% 3,30-diaminobenzidine tetrahydrochloride (DAB; Sigma, St. Louis, MO, USA) and 0.01% hydrogen peroxide in PBS, until a dark brown reaction product was evident (∼12 minutes). Sections were rinsed and mounted on glass slides, air dried, stained with cresyl violet and coverslipped.

### Cell Counting Quantification

Acute slices containing the dorsal dentate gyrus from the injected hemisphere of male (n = 2) and female (n = 2) C57Bl/6J mice injected with both AAV.mscRE4.YFP and AAV.CaMKII.mCherry were imaged on a Nikon C2 laser-scanning confocal system with an inverted Nikon ECLIPSE Ti2 microscope (10x objective, dry). Cells were counted in the hilus and the granule cell layer from two acute slices per animal. Cell counting was performed using the freehand selection tool and ROI manager in ImageJ/FIJI. Images were first assessed looking at the TRITC channel alone, so putative cells could first be confirmed as mCherry+ (CaMKII-expressing) before assessing the YFP signal. If the fluorescence value of a putative cell in the TRITC channel exceeded that of the mean plus three times the standard deviation of the background signal in that image, then the cell was considered mCherry+. If a putative cell was mCherry+, we then changed the image to the FITC channel to measure the same cell for YFP signal (mscRE4 expression). If the fluorescence value of YFP in that cell exceeded the mean plus three times the standard deviation of the background in the FITC channel, that cell would also be considered YFP+. All putative cells in the hilar region were counted, while in the granule cell layer 45 cells per slice were counted (N = 360). Raw data was analyzed in Excel to determine the percentages of mCherry+/YFP+ and mCherry+/YFP-cells in the hilar region and granule cell layer of each acute slice. These percentages were then inputted into GraphPad Prism for figure generation.

### Line Scan Profiles in the DG

Acute slices containing the dorsal dentate gyrus from both the injected and non-injected hemisphere of co-injected AAV.mscRE4.YFP/AAV.CaMKII.mCherry male (n = 1) and female (n = 2) C57Bl/6J mice were imaged on a Nikon C2 laser-scanning confocal system with an inverted Nikon Ti2 microscope (10x objective, dry). Three acute slices from each the injected and non-injected hemispheres were analyzed per animal. Line scan profiles were performed using the straight-line tool and ‘Plot Profile’ analysis in ImageJ/FIJI. Ten lines were drawn from the innermost border of the granule cell layer to the edge of the outer molecular layer on the DAPI channel of each image. Lines were drawn approximately equidistant from one another, evenly spanning the upper and lower blades of the dentate gyrus. Line scan values were then obtained on each the DAPI, mCherry, and YFP channels. As the thickness of the hippocampus varies along its axis, fluorescence values from each line scan were fed into a custom written algorithm, which normalized all values to the mean and subsequently interpolated fluorescence values based on a standardized distance (0-1). This data was then inputted into GraphPad Prism for figure generation.

---

**Supplemental Figure 1.**
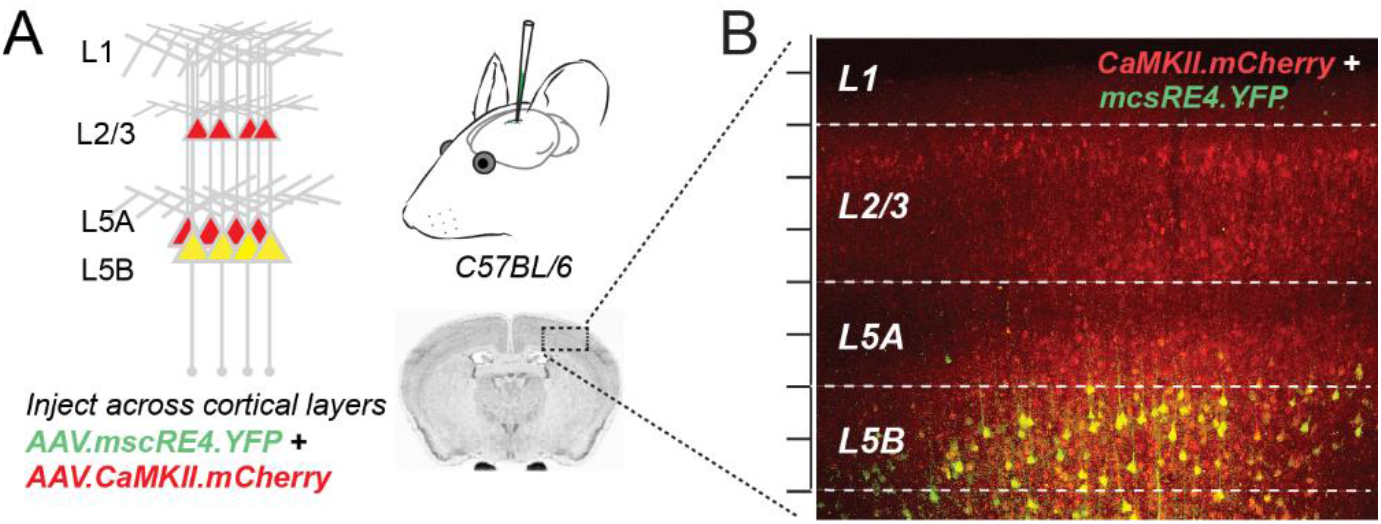
AAV.mscRE4.YFP labels layer 5B pyramidal neurons in primary motor cortex. (A) AAV.mscRE4.YFP and AAV.CaMKII.mCherry were co-injected into one hemisphere of the primary motor cortex (M1) in adult wild-type mice. (B) AAV.mscRE4.YFP expression was directly mainly in layer 5B pyramidal neurons in M1, as indicated by co-expression of AAV.mscRE4.YFP (green) and AAV.CaMKII.mCherry (red). Scale between hash marks: 50 µm.

**Supplemental Figure 2.**
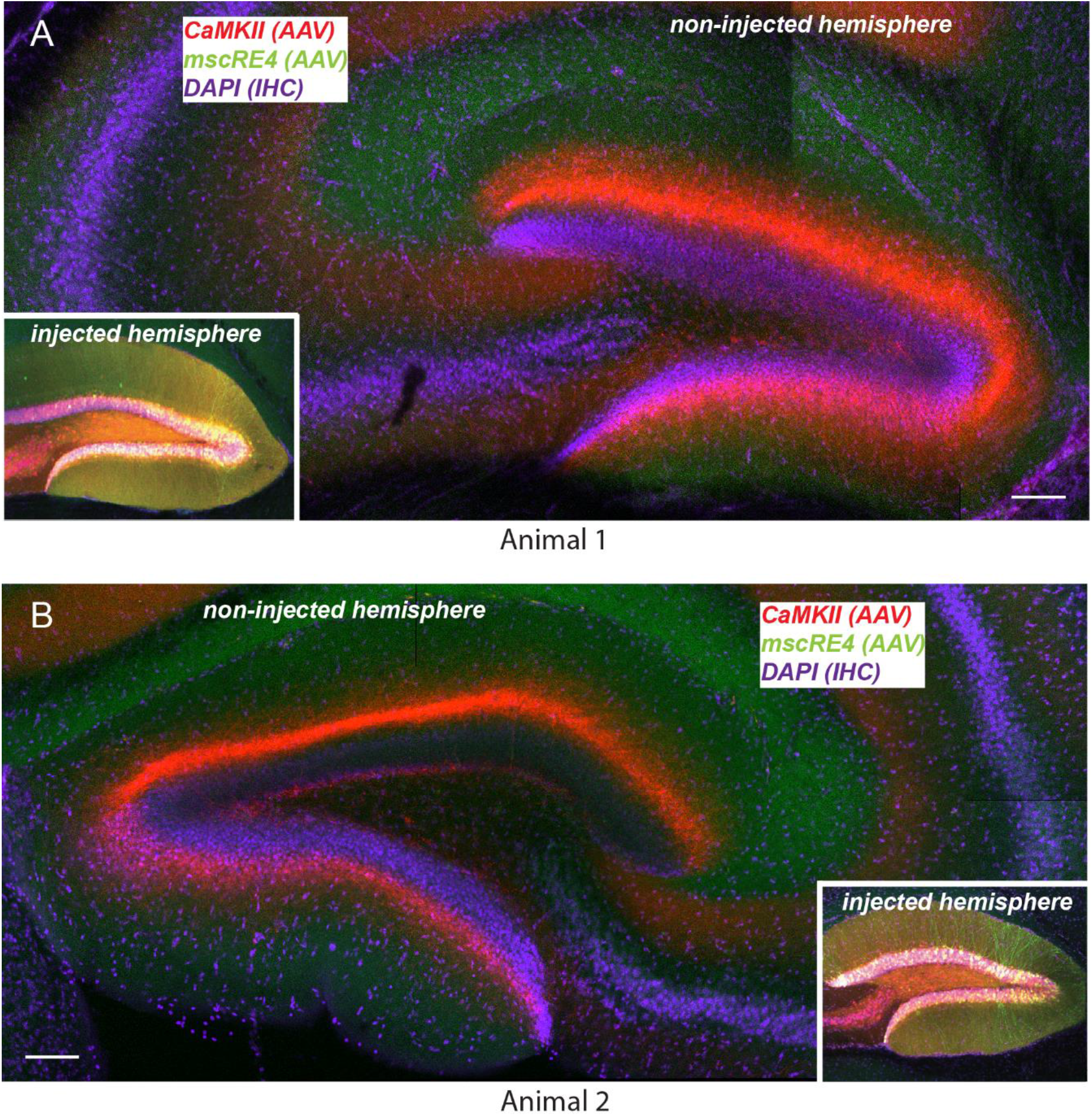
AAV.mscRE4.YFP and AAV.CaMKII.mCherry labeling in non-injected DG hemispheres. (A, B) Representative confocal images (10x objective) of the non-injected dentate gyrus from two wild-type mice injected with AAV.mscRE4.YFP and AAV.CaMKII.mCherry in the contralateral dentate gyrus (insets). Scale bars: 100 µm.

**Supplemental Figure 3.**
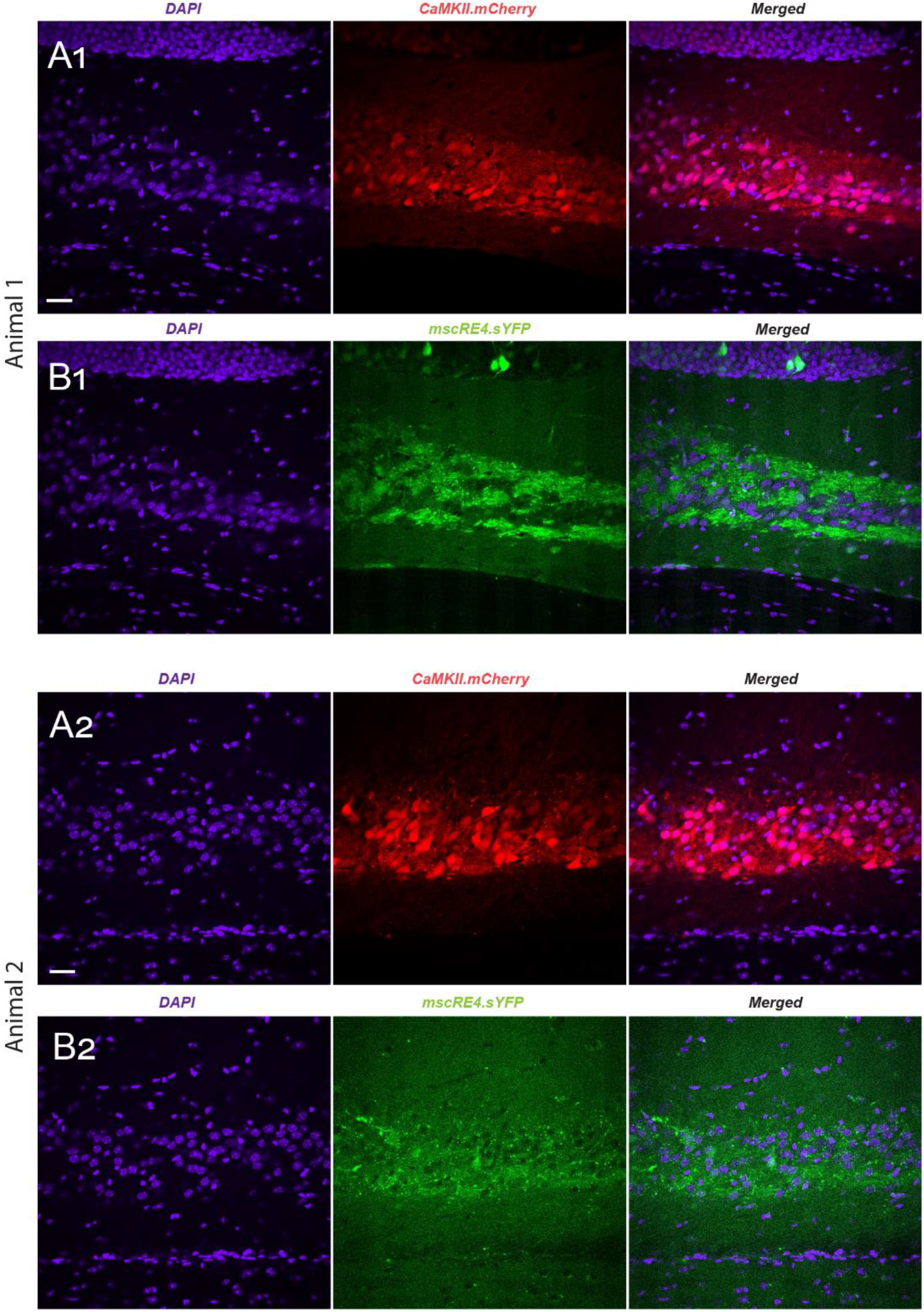
AAV.mscRE4.YFP expression in mossy fibers but not in CA3 pyramidal neurons. (A1, A2) Representative confocal images (40x objective, oil) of the CA3 region from two wild-type mice injected in one hemisphere dentate gyrus with AAV.mscRE4.YFP and AAV.CaMKII.mCherry, notable for the overlap between the DAPI (blue) and AAV.CaMKII.mCherry (red) signals compared to (B1, B2) the lack of overlap between the signals for DAPI and AAV.mscRE4.YFP (green). Scale bars: 50 µm.

**Supplemental Figure 4.**
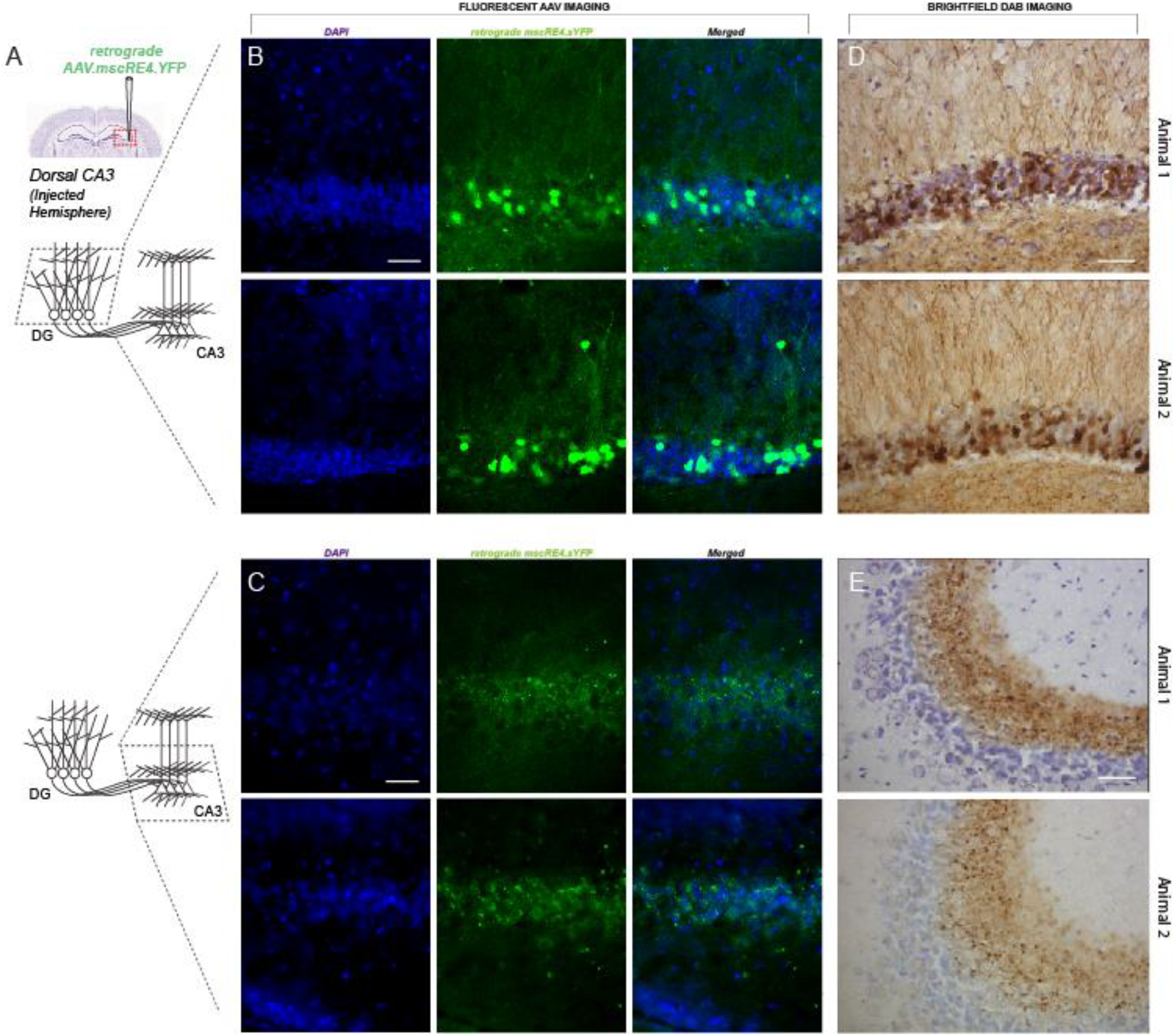
AAV.retro.mscRE4.YFP produces a similar expression pattern to AAV.mscRE4.YFP. (A) The YFP-expressing mscRE4 construct was packaged into a retrograde capsid and injected at the CA3b/CA3a border in adult wild-type mice. After 1-2 weeks, fluorescent images (40x objective) (B) in the DG showed YFP expression in granular layer cell somas and dendrites in the molecular layer, while (C) CA3 showed punctate YFP expression surrounding, but not overlapping with, stratum pyramidale cell somas. Brightfield images (40x objective) of slices which received immunostaining against GFP show abundant dark brown reaction products in (D) DG granular layer somas and molecular layer dendrites, as well as (E) punctate expression in CA3. Scale bars: 50 µm.

**Supplemental Figure 5.**
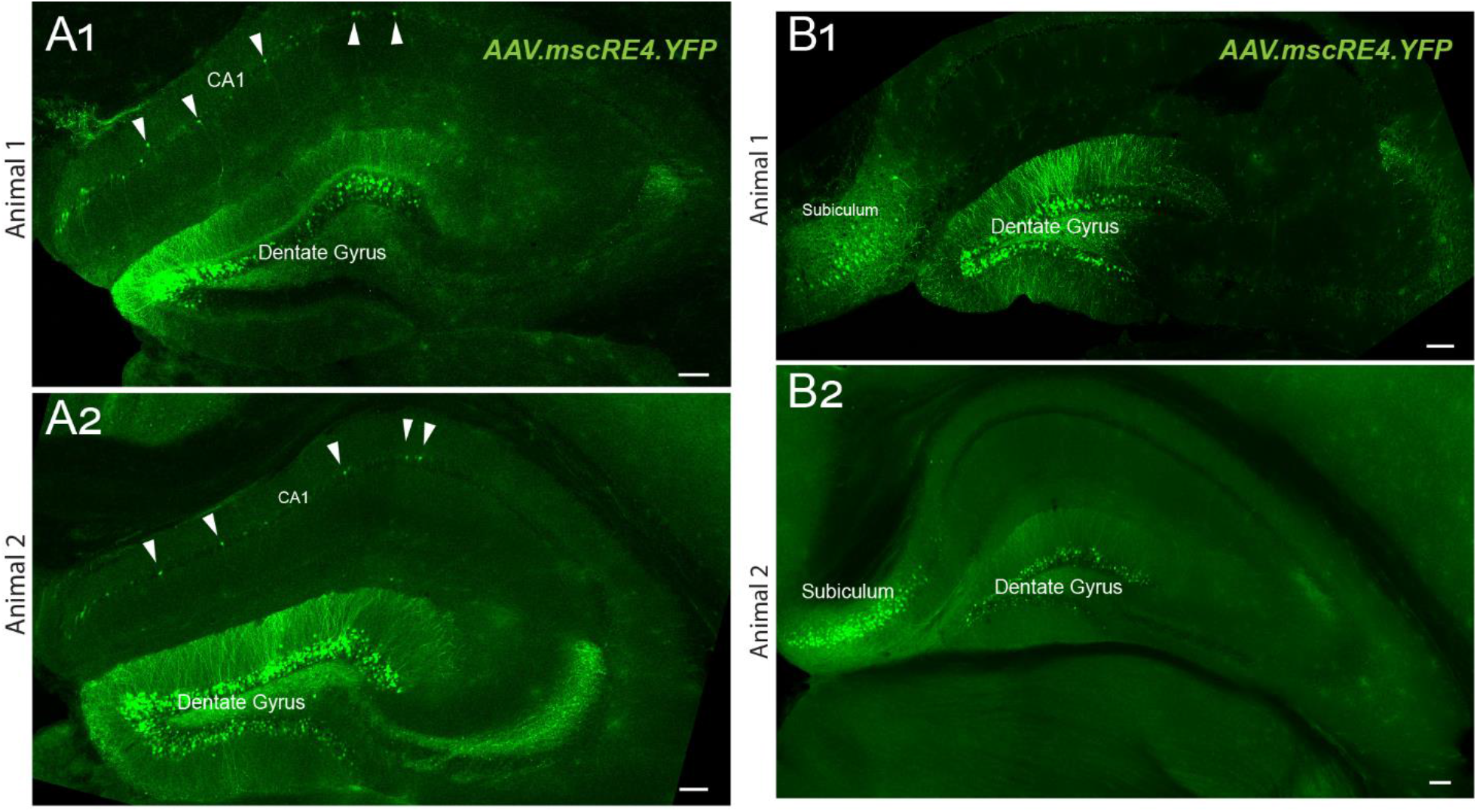
AAV.mscRE4.YFP expression in CA1 and the subiculum. (A1, A2) Representative confocal images (10x objective, dry) from two wild-type mice injected in the dentate gyrus in both hemispheres with AAV.mscRE4.YFP, showing sparse AAV.mscRE4.YFP expression in putative pyramidal neurons (white arrows) in the CA1 region, as well as (B1, B2) dense AAV.mscRE4.YFP expression in the subiculum. Scale bars: 100 µm.

